# Co-culturing *Hyphomicrobium nitrativorans* strain NL23 and *Methylophaga nitratireducenticrescens* strain JAM1 allows sustainable denitrifying activities under marine conditions

**DOI:** 10.1101/2021.02.10.430622

**Authors:** Cucaita Alexandra, Piochon Marianne, Villemur Richard

## Abstract

*Hyphomicrobium nitrativorans* strain NL23 and *Methylophaga nitratireducenticrescens* strain JAM1 were the principal bacteria involved in the denitrifying activities of a methanol-fed, fluidized marine denitrification reactor. We believe that a tight relationship has developed between these two strains to achieve denitrification in the reactor under marine conditions. To characterize the potential synergy between strain JAM1 and strain NL23, we compared some of their physiological traits, and performed co-cultures. Pure cultures of strain JAM1 had a readiness to reduce nitrate (NO_3_−) with no lag phase for growth contrary to pure cultures of strain NL23, which has a 2-3 days lag phase before NO_3_− starts to be consumed and growth to occur. Compared to strain NL23, strain JAM1 has a higher μmax for growth and higher specific NO_3_− reduction rates. Antagonist assays showed no sign of exclusion by both strains. Planktonic co-cultures could only be performed on low NaCl concentrations for strain NL23 to survive. Denitrification rates were twice higher in the planktonic co-cultures than those measured in strain NL23 pure cultures. Biofilm co-cultures were performed for several months in a 500-mL bioreactor filled with Bioflow supports, and operated under fed-batch mode with increasing concentrations of NaCl for strain NL23 to acclimate to marine conditions. Under these conditions, the biofilm co-cultures showed sustained denitrifying activities and surface colonization by both strains. Increase in ectoine concentrations produced by strain JAM1 was observed in the biofilm with increasing NaCl concentrations. These results illustrate the capacity of both strains to act together in performing denitrification under marine environments. Although strain JAM1 did not contribute in better specific denitrifying activities in the biofilm co-cultures, its presence was essential for strain NL23 to survive in a medium with NaCl concentrations > 1.0%. We believe that ectoine is an important factor for the survival of strain NL23 in these environments.

## INTRODUCTION

Denitrification takes place in bacterial cells where N oxides serve as terminal electron acceptor instead of oxygen (O_2_) for energy production when O_2_ depletion occurs, leading to the production of gaseous nitrogen (N_2_). Four sequential reactions are required for the reduction of nitrate (NO_3_−) to gaseous nitrogen, via nitrite (NO_2_−), nitric oxide and nitrous oxide, and each of these reactions is catalyzed by different enzymes, namely NO_3_− reductases (Nar or Nap), NO_2_− reductases (Nir), nitric oxide reductases (Nor) and nitrous oxide reductases (Nos) [1–3].

We have been studying for many years the biofilm derived from a continuous, fluidized-bed methanol-fed denitrification reactor that treated a 3 million-L seawater aquarium that contains fish, birds and invertebrates. The fluidized carriers in the denitrification reactor were colonized by naturally occurring multispecies bacteria to generate a marine methylotrophic denitrifying biofilm [4, 5], among which the methylotrophic bacteria *Methylophaga* spp. and *Hyphomicrobium* spp. accounted for 60 to 80% of the bacterial community [6]. Two bacterial strains representative of *Methylophaga* spp. and *Hyphomicrobium* spp. were isolated from the denitrifying biofilm. The first one, *Methylophaga nitratireducenticrescens* strain JAM1, is capable of growing in pure cultures under anoxic conditions by reducing NO_3_− to NO_2_−, which accumulates in the medium [7, 8]. It was later shown to be able to reduce NO and N_2_O to N_2_ [9]. These activities concur with the presence of gene clusters encoding two Nar reductases, two Nor reductases and one Nos reductase [10, 11]. A dissimilatory NO-forming NO_2_− reductase gene (*nirS* or *nirK*) is absent, which correlates with accumulation of NO_2_− in the culture medium during NO_3_− reduction. The second strain, *Hyphomicrobium nitrativorans* strain NL23, is capable of complete denitrification from NO_3_− to N_2_, and possesses operons that encode for the four different nitrogen oxide reductases, among which a periplasmic Nap-type NO_3_− reductase [12–14]. These two species are the main bacteria responsible of the dynamics of denitrification in the biofilm.

*Hyphomicrobium* spp. are methylotrophic bacteria that are ubiquitous in the environment [15]. They have also been found in significant levels in several methanol-fed denitrification systems treating municipal or industrial wastewaters or a seawater aquarium, and they occurred often with other denitrifying bacteria such as *Paracoccus* spp., *Methylophilales* or *Methyloversatilis* spp. Their presence correlates with optimal performance of bioprocess denitrifying activities [16–22]. *Methylophaga* spp. are methylotrophic bacteria isolated from saline environments [23, 24]. They have been found in association with diatoms, phytoplankton blooms and marine algae, which are known to generate C1 carbons [25–27]. Co-occurrence of *Methylophaga* spp. and *Hyphomicrobium* spp. has been shown in two other methanol-fed denitrification systems treating saline effluents [28, 29]. Therefore, understanding how these two taxa collaborate could benefit in optimizing denitrification systems treating saline/brackish waters.

We hypothesize that a tight relationship has developed between *H. nitrativorans* strain NL23 and *M. nitratireducenticrescens* strain JAM1 to achieve denitrification in the original biofilm. For instance, because growth of strain NL23 is impaired in marine environments [12], strain JAM1 would protect strain NL23 from osmotic shock in the biofilm by producing, for instance, the osmoprotectant ectoine, as most of *Methylophaga* species can generate [23]. In the present report, we compared some physiological traits of both strains that can influence their co-habitation. We then used co-culture approach to characterize the potential synergy between these two strains to achieve denitrification. Our results showed that when both strains are present, sustainable denitrifying activities were achieved and that strain JAM1 is essential for strain NL23 to survive under marine conditions.

## MATERIAL AND METHODS

### Determination of NO_3_−, NO_2_− and biomass

NO_3_− and NO_2_− were measured either by ionic chromatography as described by Mauffrey *et al.* [9] or by colorimetry based on the method described by Schnetger and Lehners [30]. In this method, NO_3_− was reduced to NO_2_− with vanadium (III) chloride (VCl_3_). Eight hundred mg VCl_3_ were dissolved in 100 mL of 0.1 M HCl (Solution A). The solution was filtered (0.45 μm). Solution B was made of 200 mg N-1-naphthylethylenediamine dihydrochloride in 100 mL H_2_O. Solution C was made of 2 g sulfanilamide in 100 mL HCl 10%. These solutions were kept in amber bottle at 4°C for one month. Solutions D and E were made fresh. For solution D, solutions A, B and C were mixed at the proportion of 5:1:1. For solution E, solutions B and C were mixed at 1:1 proportion. The assays were carried out in 96-well plates. To determine the NO_x_ concentrations (NO_3_− + NO_2_−), samples (120 μL) were added to one plate, then 100-μL solution D was quickly added and mixed with a multichannel micropipet. The plate was covered and immediately incubated for 60 min at 45°C. To measure NO_2_− already present in the medium, the same samples (200 μL) were added to another plate, then 20-μL solution E was quickly added and mixed with a multichannel micropipet. The plate was covered and immediately incubated for 30 min at 45°C. Both plates were read at 540 nm with a plate reader. The concentrations were determined with standard solutions. Linear response ranged from 0.1 to 2 mg-N/L (either NO_3_− or NO_2_−). Results from the NO_3_− reduction by VCl_3_ generated the NO_x_ concentrations (NO_3_− + NO_2_−). Therefore, NO_3_− concentrations were calculated as NO_x_ – NO_2_−.

The growth rates, the NO_3_− reduction rates and the denitrification rates were calculated by the linear portion of the culture growth (OD_600nm_), of the NO_3_− concentrations and of the NO_x_ concentrations, respectively, over time for each replicate (OD h^−1^; NO_3_− mM h^−1^, NO_x_ mM h^−1^). The specific denitrification rates (NO_x_ mM h^−1^ mg-protein^−1^) were reported as the denitrification rates divided by the quantity of biomass (mg protein) in a vial or in the reactor. The amount of protein in the biomass was determined by the Quick StartTM Bradford Protein Assay (BioRad, Mississauga, ON, Canada).

### Culture media

The Instant Ocean (IO) seawater medium was bought from Aquarium systems (Mentor, OH, USA) and dissolved at 30 g/L. NaNO_3_ was added at prescribed concentrations. The medium was adjusted at pH 7.5 and autoclaved. One ml per liter of autoclaved trace metal solution (per liter: 0.9 g FeSO_4_.7H_2_O, 0.03 g CuSO_4_.5H_2_O, 0.234 g MnCl_2_.4H_2_O and 0.363 g Na_2_MoO_4_.2H_2_O) was added. Three mL of 0.2 μm-filtered methanol were added per liter of medium.

Planktonic pure cultures of *H. nitrativorans* NL23 were performed in the 337a medium (per liter: 1.3 g KH_2_PO_4_, 1.13 g Na_2_HPO_4_, 0.5 g (NH_4_)_2_SO_4_, 0.2 g MgSO_4_.7H_2_O). NaNO_3_ was added at the prescribed concentrations. The medium was adjusted at pH 7.5 and autoclaved. These 0.2 μm-filtered solutions were added per liter: 5 mL trace element solution (per liter: 309 mg CaCl_2_.2H_2_O, 200 mg FeSO_4_.7H_2_O, 100 mg Na_2_MoO_4_.2H_2_O, 67 mg MnSO_4_.H_2_O), 3 mL methanol and 1mL vitamin B_12_ (0.1 mg/ml).

Planktonic pure cultures of *M. nitratireducenticrescens* JAM1 were performed in the *Methylophaga* 1403 medium (per liter: 24 g NaCl, 3 g MgCl_2_. 6 H_2_O, 2 g MgSO_4_.7H_2_O, 0.5 g KCl, 1 g CaCl_2_, 0.5 g Bis-tris, and NaNO_3_ at the prescribed concentrations). The medium was autoclaved, and these 0.2 μm-filtered solutions were added per liter: 3 mL methanol, 20 mL solution T (per 100 mL: 0.7 g KH_2_PO_4_, 10 g NH_4_Cl, 10 g Bis-tris, 0.3 g citrate ferric ammonium, pH 8.0), 1 mL vitamin B_12_ (0.1 mg/ml), 10 ml Wolf solution [per liter: 0.5 g EDTA, 3.0 g MgSO_4_.7H_2_O, 0.5 g MnSO_4_.H_2_O, 1.0 g NaCl, 0.1 g FeSO_4_.7H_2_O, 0.1 g Co(NO_3_)_2_.6H_2_O, 0.1 g CaCl_2_ (anhydrous), 0.1 g ZnSO_4_.7H_2_O, 0.010 g CuSO_4_.5H2O, 0.010 g AlK(SO_4_)_2_ (anhydrous), 0.010 g H_3_BO_3_, 0.010 g Na_2_MoO_4_.2H_2_O, 0.001 g Na_2_SeO_3_ (anhydrous), 0.010 g Na_2_WO_4_.2H_2_O, and 0.020 g NiCl_2_.6H_2_O, pH 8.0]. For some assays, the NaCl concentration was adjusted in this medium as needed (from 0% to 2.75%). For solid media, agar (1.5% final concentration) was added before sterilization.

### Planktonic pure cultures and co-cultures

Generation of pre-cultures were derived from cultures of strain JAM1 and strain NL23 cultured under oxic conditions without NO_3_− (unless specified) in their respective medium (*Methylophaga* 1403 and 337a) and incubated at 30°C at 150 rpm. The cells were then centrifuged and dispersed in the prescribed medium before use. For anoxic cultures, the sterile media were distributed (60 mL) in sterile serologic vials. The vials were purged of oxygen for 10 min with nitrogen gas (Praxair, Mississauga, ON, Canada) and sealed with sterile septum caps. The planktonic co-cultures (anoxic conditions) were inoculated with pre-cultures of strain JAM1 and of strain NL23 with a JAM1/NL23 ratio of 1:10 and a final cell dispersion of 0.1 OD_600nm_. Cultures were incubated for 1-7 days at 30°C without shaking.

### Biofilm co-cultures in recirculating reactor

The reactor is illustrated in Figure 1. It consisted of 500-ml working volume. A flow cell consisted of a 50-mL tube was added in the recirculating circuit where two microscope slides were added to monitor cell attachment on the surface. The system was carefully washed with water and ethanol. Two hundred and sixty acid-washed Bioflow 9 mm supports were added in the reactor. The media (Table 1) were sterilized before used, but was added not sterilely under oxic environment. Throughout all the different operating conditions, the medium was recirculated with a peristaltic pump at 30 mL/min, and the reactor was run at room temperature (*ca.* 22°C). Gas production was recorded by water displacement in a graduated cylinder (Fig. 1). Samples of the suspended biomass (1 mL) were taken at sampling ports and centrifuged; the supernatant was used to determine the NO_3_− and NO_2_− concentrations. The pellets of some of these samples were kept for DNA or protein (or both) extractions. From the DNA extracts, the concentrations of strain JAM1 and strain NL23 in the suspended biomass were determined by qPCR.

**Table 1.**
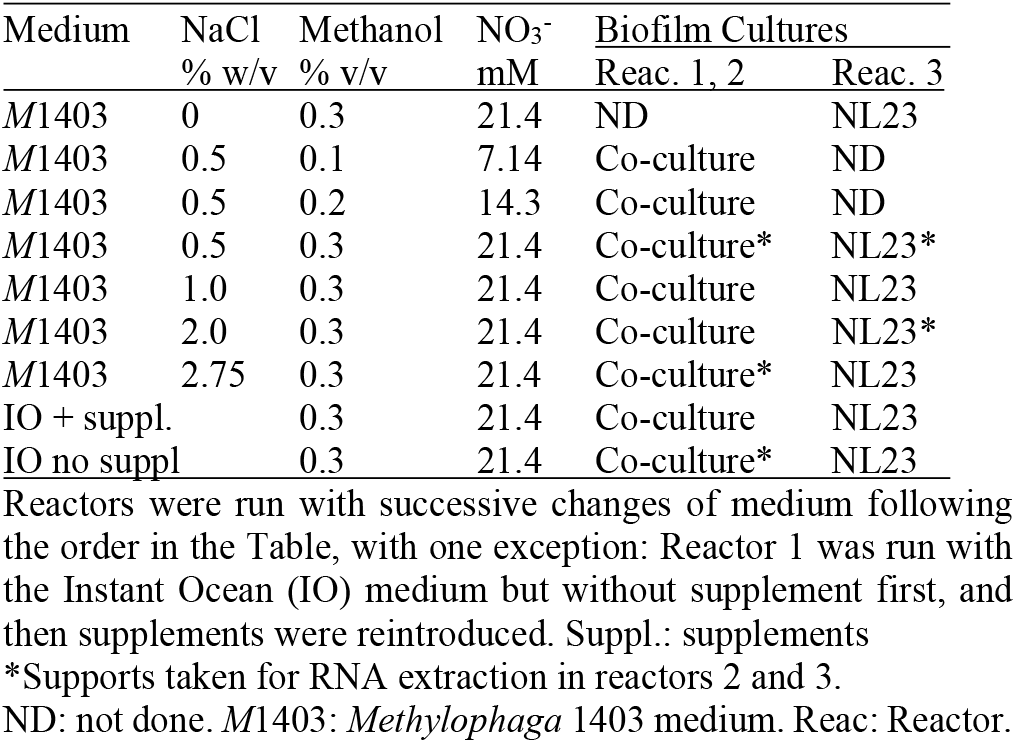
Operating conditions of the reactors.

| Medium | NaCl<br>% w/v | Methanol<br>% v/v | NO<sub>3</sub><sup>-</sup><br>mM | Biofilm Cultures |  |
| --- | --- | --- | --- | --- | --- |
|  |  |  |  | Reac. 1, 2 | Reac. 3 |
| M1403 | 0 | 0.3 | 21.4 | ND | NL23 |
| M1403 | 0.5 | 0.1 | 7.14 | Co-culture | ND |
| M1403 | 0.5 | 0.2 | 14.3 | Co-culture | ND |
| M1403 | 0.5 | 0.3 | 21.4 | Co-culture* | NL23* |
| M1403 | 1.0 | 0.3 | 21.4 | Co-culture | NL23 |
| M1403 | 2.0 | 0.3 | 21.4 | Co-culture | NL23* |
| M1403 | 2.75 | 0.3 | 21.4 | Co-culture* | NL23 |
| IO + suppl. |  | 0.3 | 21.4 | Co-culture | NL23 |
| IO no suppl |  | 0.3 | 21.4 | Co-culture* | NL23 |

Reactors were run with successive changes of medium following the order in the Table, with one exception: Reactor 1 was run with the Instant Ocean (IO) medium but without supplement first, and then supplements were reintroduced. Suppl.: supplements
\*Supports taken for RNA extraction in reactors 2 and 3.
ND: not done. M1403: *Methylophaga* 1403 medium. Reac: Reactor.

**Figure 1.**
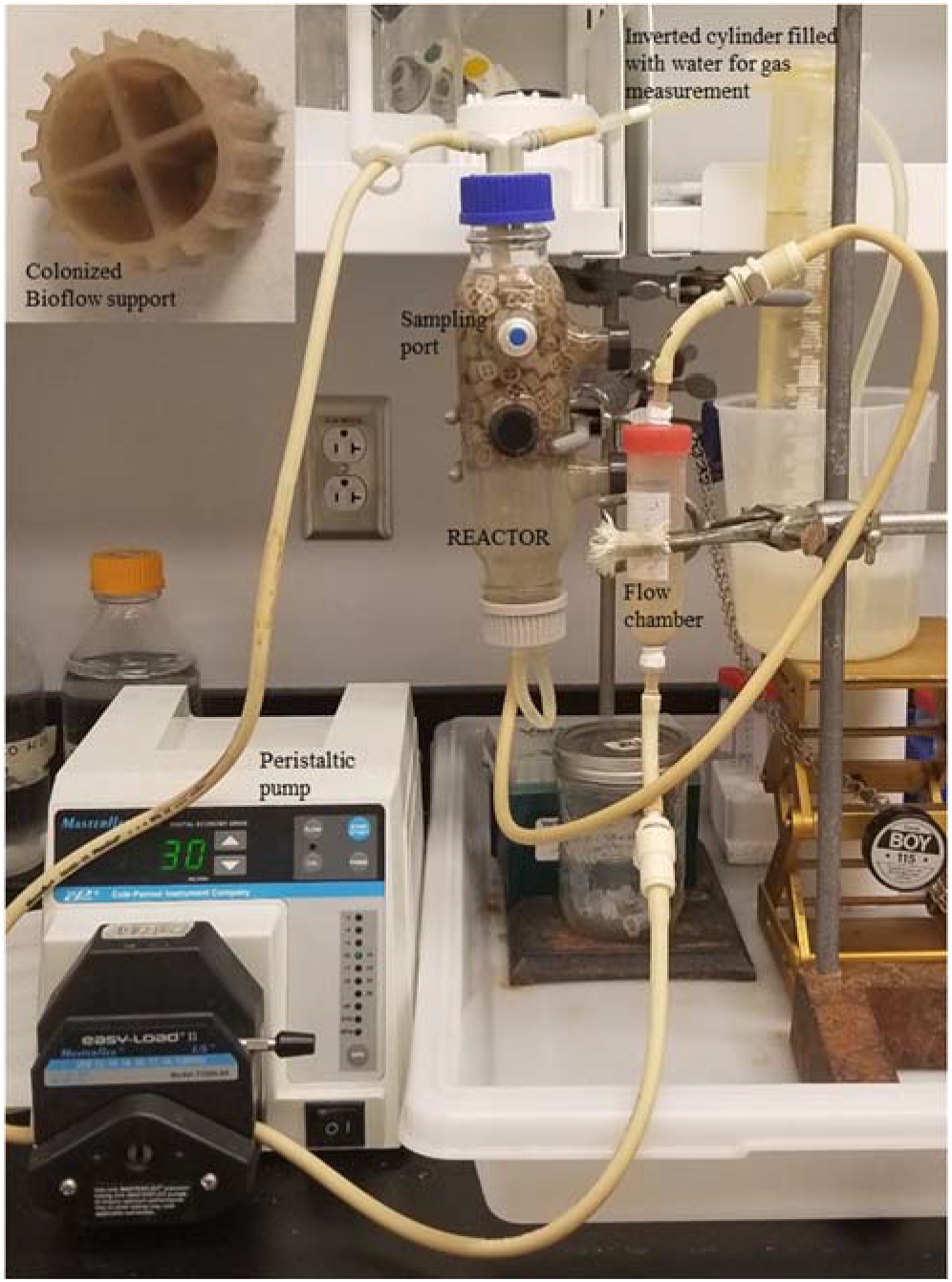
Configuration of the reactor.

The same reactor was used to carry out three runs sequentially. Reactors 1 and 2 were inoculated with strain JAM1 and strain NL23 (biofilm co-cultures). Reactor 3 was inoculated with only strain NL23 (biofilm mono-culture). Results from reactor 1 allowed to adjust the conditions for reactor 2. In reactors 2 and 3, protein and RNA extractions, and microscopic observations were performed; ectoine concentration was measured in the biofilm of reactor 2.

To initiate reactors 1 and 2 (co-cultures), inocula of strain NL23 (four 60 mL-vials) and strain JAM1 (three 60 mL-vials) were centrifuged (*ca.* 3 g wet weight, each), dispersed in 20 mL 0.5% NaCl *Methylophaga* 1403 medium, and added to the reactor filled with 500 mL of the same medium without NO_3_−. To acclimate both strains to the reactor environment, the oxic conditions were kept by injecting air with a peristaltic pump for 3 days in the reactor, after which air was no longer supplied and NO_3_− was added to create anoxic conditions. For reactor 3, the inoculum of strain NL23 was added in the reactor with 0% NaCl *Methylophaga* 1403 medium, and the reactor was operated directly under anoxic conditions (no oxic phase). The reactors were then run for a week, with additions of NO_3_− when needed, after which the medium was replaced by a fresh one. Only bacteria attached to supports and vessel were kept in the reactor. The planktonic bacteria were discarded. Upon NO_3_− and NO_2_− reduction, the medium was changed with fresh one. Anoxic conditions and denitrifying activities were occurring as indicated by gas production and the reduction of NO_3_− and NO_2_−. When these production and reduction appeared stable, the reactor was sampled at regular intervals for at least 24 h to derive the dynamics of NO_3_− and NO_2_− reduction. Afterwards, one to two supports were collected, frozen at −20°C until use for the extractions of proteins, DNA and ectoine. In some of the operating conditions (Table 1), supports were collected for RNA extractions (reactors 2 and 3. To do so, fresh medium was added in the reactor. When NO_3_− was near completely consumed, 10 supports were collected in a glove box flushed with nitrogen gas and frozen immediately in liquid nitrogen, and kept at −70°C until use to extract total RNA. In all cases, the taken supports were replaced with virgin ones in the reactor. After these steps, we proceeded with new medium (increasing concentration of methanol/NO_3_− or NaCl) as described in Table 1. When the different operating runs for reactor 1 were finished, all the supports were frozen at −20°C for further analysis. The vessel was then washed thoroughly and sterilized. New virgin supports were added to start reactor 2. The same was done with reactor 3.

### Ectoine

The protocol was based on Chen et al. [31] and Zhang et al. [32]. The biofilm on one support was directly extracted with 2 ml of methanol/chloroform/water (10:5:4) with intermittent strong agitation for 3 h. The extract was then centrifuged at 16 000 *g* for 10 min. One mL of the supernatant was extracted with 1 mL chloroform/water (1:1) with agitation for 30 min. The extract was then centrifuged at 16 000 *g* for 10 min. The water phase was collected and dried at 35°C with N_2_ gas for 30 min. The extract was then dissolved in 1 mL water and filtered on 0.2 μm filter. The stable isotopically labelled internal standard 5,6,7,8-tetradeutero-4-hydroxy-2-heptylquinoline (HHQ-d_4_) was added in the samples (2.66 μg/mL final concentration). HHQ-d_4_ was provided by the laboratory of Eric Déziel (INRS, Laval, QC, Canada). Ectoine ((*S*)-2-methyl-1,4,5,6-tetrahydro-pyrimidine-4-carboxylic acid, 97% purity) was bought at Sigma Aldrich (Oakville, ON, Canada), and used to derive a standard curve.

Fifteen-μL of the samples were analysed by a HPLC-coupled to a mass spectrometer (Waters, Milford, MA, USA). The Waters 2795 HPLC system was equipped with a Kinetex (100 x 4.6 mm) 2.6-μm C8 reverse-phase LC column (Phenomenex, Torrance, CA, USA). The mobile phase was a gradient of 1% acetic acid in water (solvent A) and 1% acetic acid in acetonitrile (solvent B) programmed as follows: initial 2% solvent B (0–4 min), 2–70% solvent B (4–5 min), 100% solvent B (5–8 min), then hold 3 min and followed by 3 min of re-equilibration. HPLC flow rate was 400 μl/min split to 40 μl/min by a Valco tee splitter. The Quattro Premier XE mass spectrometer was equipped with a Z-spray interface using electro-spray ionization in positive mode (ESI-MS/MS). Multiple Reaction Monitoring (MRM) mode was used to quantify ectoine. MassLynx and QuanLynx software (Ver. 4.1) were used. The capillary voltage was set at 3.0 kV and the cone voltage at 30 V. The source temperature was kept at 120°C. Nitrogen was used as nebulising and drying gas at flow rates of 15 and 100 ml/min, respectively. In MRM mode, the following transitions were monitored: for ectoine 143→98 and for the internal standard HHQ-d4 248→163. The pressure of the collision gas (argon) was set at 2 × 10^−3^ mTorr and the collision energy at 30 V for all transitions. The area of each chromatographic peak was integrated and the ectoine/HHQ-D_4_ response ratio was used to create the calibration curve (R^2^ = 09945) and quantify the concentration of ectoine in the samples.

### Antagonist assays

Cultures of strain JAM1 or strain NL23 (each OD_600nm_ 0.5) were first spread (200 μL) over *Methylophaga* 1403 (0.5% NaCl, no NO_3_−) agar plates with 0.3% methanol. From these, three protocols were carried out. In the first protocol, one-cm^2^ blotting papers were soaked with either strain NL23 culture or strain JAM1 culture. The papers were then put on the inoculated agar plates. In the second protocol, one drop of strain JAM1 and strain NL23 cultures was put onto the inoculated agar plates. In the third protocol, a well was made in the inoculated agar plate and one drop of either strain JAM1 or strain NL23 cultures was put in it. These plates were incubated between 30 and 48 h at 30°C under oxic conditions. Clearing zone around the inoculated points or the absence of growth at these points were recorded.

### Fluorescence *in situ* hybridization (FISH)

The biofilm that colonized the microscope slides in the flow cell chamber was fixed with 4% cold paraformaldehyde in PBS for 60 min, and washed twice with water. The biofilm was dehydrated for 3 min successively in 50%, 80% and 95% ethanol/water, and then air dried. Hybridization buffer for both probes consisted of 20% formamide, 20 mM Tris-HCl pH 8.0, 0.01% SDS and 0.9 M NaCl. The probe concentrations used were between 10 to 20 ng/μL. Hybridization was carried out in an Omnislide *in situ* thermal cycler (Thermo Electron Corporation, Waltham, Mass., USA) for 3 h at 46°C, followed with by 2 min at 48°C. Slides were washed in 20 min at 48°C, in 20 mM Tris-HCl pH 8.0, 5 mM EDTA, 0.01% SDS and 210 mM NaCl. FISH samples were mounted with Prolong Gold agent containing DAPI (Molecular Probes, Fisherscientific, Ottawa, ON Canada). The probes for *Methylophaga* spp. (MPH-730-Atto488; 5’ CAGTAATGGCCCAGTGAGTCGCC 3’) [33] and for *Hyphomicrobium* spp. (Hypho-Cy3; 5’ TCCGTACCGATAGGAAGATT 3’) [34] were synthesized by Biomers.net (Ulm, Germany) and AlphaDNA (Montreal, QC, Canada), respectively. Slides were examined on an epifluorescence Leica DM3000 microscope.

### DNA and RNA extraction, qPCR and RT-qPCR, and RNA sequencing

DNA and RNA extractions of the planktonic biomass and of the biofilm from 1 to 4 supports were performed as described [5, 9, 35]. Concentrations of strain JAM1 and strain NL23 were performed by qPCR with primers targeting *narG1* and *napA*, respectively as described by Geoffroy *et al*. [10] (Table 2). Standard curves were performed with PCR-amplified DNA fragments of *narG1* and *napA* from strain JAM1 genome and strain NL23 genome, respectively (Table 2). RT-qPCR were performed as described by Mauffrey *et al.* with primers targeting *narG1* and *narG2* for strain JAM1, *napA* and *nirK* for strain NL23, and the reference genes *rpoB* and *dnaG* specific for each strain (Table 2). The standard curves were performed with dilution of the respective genomes.

**Table 2.**
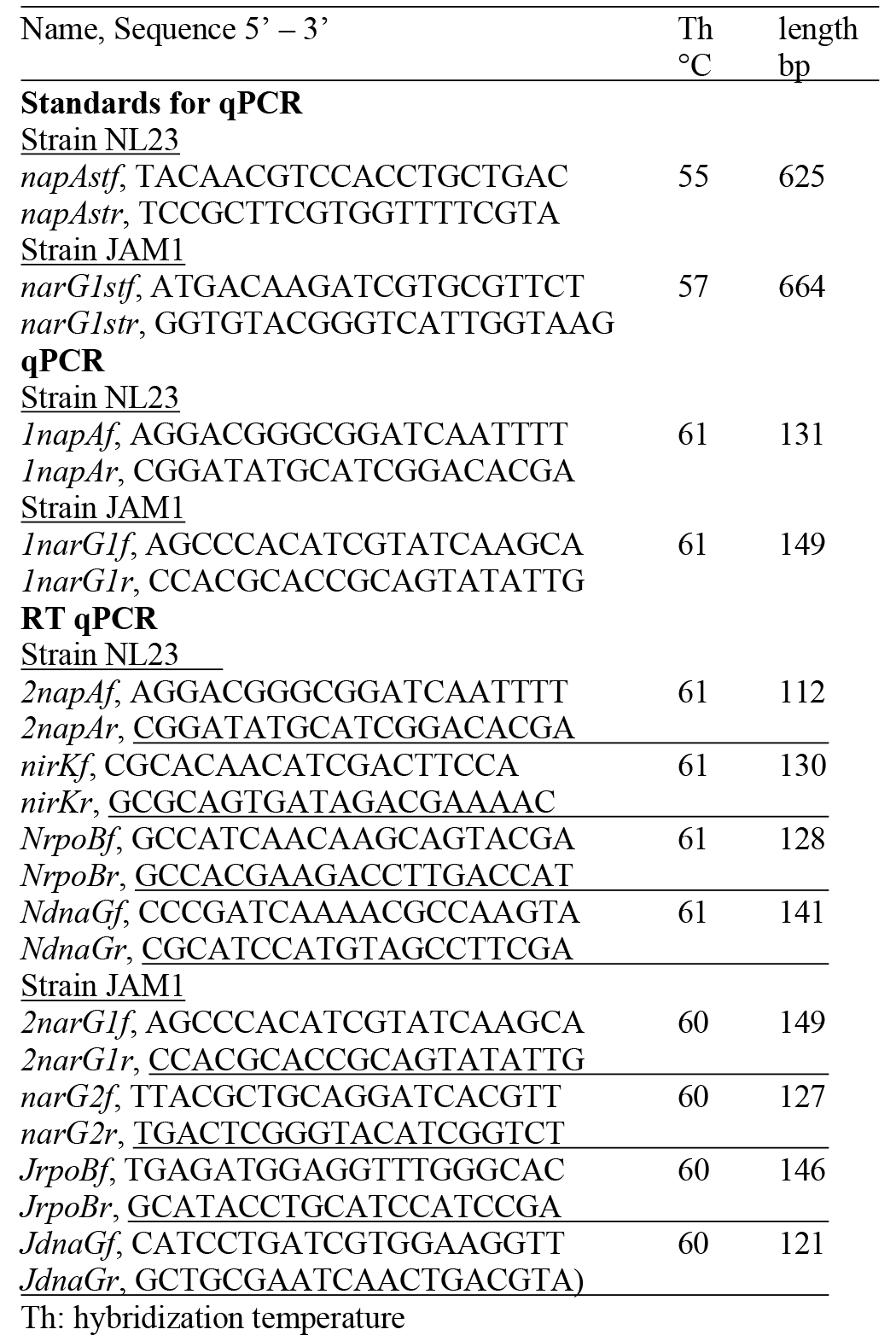
Sequences of the oligonucleotides used for qPCR and RT-qPCR assays.

The RNA samples were sequenced at the Centre d’expertise et de services Génome Québec Montreal QC, (Canada) (Sequencing type: Illumina NovaSeq 6000 S4 PE100 - 25M reads; Library Type: rRNA-depleted for bacteria). The sequencing data were uploaded to the Galaxy web platform, and we used the public server at usegalaxy.org to analyze the data [36]. Raw reads were filtered to remove low quality reads using FASTX toolkit by discarding any reads with more than 10% nucleotides with a PHRED score <20. The resulting reads were aligned to the genome of *M. nitratireducenticrescens* strain JAM1 (GenBank accession number CP003390.3) and to the genome of *H. nitrativorans* strain NL23 (CP006912.1) using Bowtie2 with default parameters. BEDtools were used to assign the number of reads to the respective genes in the genome, which were then normalized as transcripts per million (TPM). Because the samples were from one reactor run, statistical analysis could not have been performed. However, an arbitrary variation of 10% in the read values revealed that the ratio of TPM of a gene from one condition to another of >2 or <0.2 is significantly different.

## RESULTS

### Physiological differences between the two strains

We measured the dynamics of NO_3_− uptake by *M. nitratireducenticrescens* strain JAM1 and *H. nitrativorans* strain NL23. Planktonic anoxic pure cultures of strain JAM1 and strain NL23 were performed with different concentrations of NO_3_− to derive their respective maximum growth rates (μmax) and half-saturation constants of NO_3_− concentration for growth (Table 3). Strain NL23 cultures showed a 48 to 72-h lag before growth occurred, whereas strain JAM1 cultures presented no such lag (Fig. 2A). The μmax of strain JAM1 cultures were 44% higher than those of strain NL23 cultures (Fig. 2B, Table 3). To assess the affinity of these strains toward NO_3_− for growth, the μmax/Ks ratio was calculated [37] (Table 3). This ratio in strain JAM1 cultures was twice higher than that of strain NL23 cultures. The NO_3_− reduction rates in the NL23 cultures increased linearly with the increase of NO_3_− concentrations (from 2 to 40 mM) in the medium (Fig. 2C). These rates reached a plateau at 24 mM NO_3_− in strain JAM1 cultures, and showed no significant changes at higher concentrations. The specific NO_3_− reduction rates (rates normalized by the biomass) were however, relatively constant for both strains whatever the NO_3_− concentrations (Fig. 2D). These specific rates were six times higher in strain JAM1 cultures (average 4.5 ± 0.8 NO_3_− h^−1^ OD^−1^) than in strain NL23 cultures (average 0.77 ± 0.32 NO_3_− h^−1^ OD^−1^). All these results suggest that strain JAM1 has a higher dynamism towards NO_3_− than strain NL23.

**Table 3.** Kinetics of growth of strains JAM1 and NL23 under anoxic conditions.

|  | Strain JAM1<sup>a</sup> | Strain NL23 |
| --- | --- | --- |
| Lag (h) | <6 | 48-72 |
| μ<sub>max</sub> (OD<sub>600nm</sub> h<sup>-1</sup>) | 0.0111 (0.0006) | 0.0077 (0.0006) |
| K<sub>s</sub> (mM) | 13.4 (2.0) | 19.2 (3.7) |
| μ<sub>max</sub>/K<sub>s</sub> (μM<sup>-1</sup> h<sup>-1</sup>) | 0.83 | 0.40 |

<sup>a</sup> Data from Mauffrey *et al.* [35] and from new measurements.

**Figure 2.**
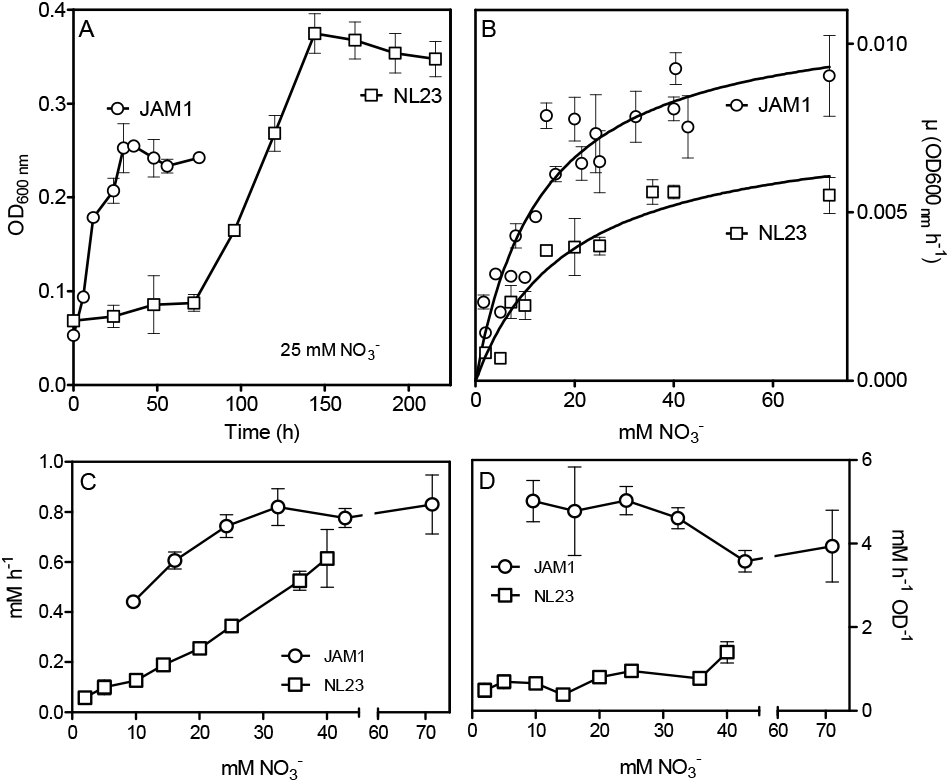
Kinetics of growth and NO_3_− reduction in planktonic mono-cultures. A. Growth under planktonic anoxic conditions (triplicate cultures). B and C. Growth rates and NO_3_− reduction rates, respectively, at different NO_3_− concentrations. Each point is the average of triplicate anoxic cultures. D. Specific NO_3_− reduction rates. These rates were calculated with the NO_3_− reduction rates by the generated culture biomass (OD_600nm_) at the end of the exponential phase. Data for strain JAM1 in panels C and D were taken from Geoffroy *et al.* [10].

The second physiological character that we tested was the capacity of both strains to produce antagonist molecules to exclude one to each other. Under the tested conditions (see M&M section), no sign of exclusion was observed from both strains.

### Anoxic planktonic co-cultures

Co-cultures were first attempted under planktonic conditions to assess the effect of culturing both strains together on the denitrifying activities. Initially, we wanted to perform planktonic co-cultures under marine conditions as both strains originated from a marine denitrification system. However, strain NL23 planktonic growth is impaired when the culture medium > 1% NaCl [12], impeding planktonic co-cultures under marine conditions. We found that the *Methylophaga* culture medium (*Methylophaga* 1403) with 0.5% NaCl instead of 2.4% can sustain both strains with minimal growth defect. This medium was used to perform our anoxic planktonic co-culture assays. Higher levels of strain NL23 in the inoculum (JAM1/NL23 ratio of 1:10) had to be used because of the higher dynamics of strain JAM1 towards NO_3_− as shown above. In the first co-culture attempts, both inocula were derived from oxic pre-cultures. Anoxic planktonic mono-cultures of both strains were performed in parallel also in this medium as control. Our results showed that these co-cultures failed to perform complete denitrification. The NO_3_− reduction rates were similar between the co-cultures (0.675 ± 0.008 mM-NO_3_− h^−1^) and strain JAM1 mono-cultures (0.815 ± 0.024 mM-NO_3_− h^−1^), and NO_2_− accumulated in the medium (Fig. 3A). The proportion of strain NL23 at the end of the co-cultures was 1.6% (Table 4).

**Table 4.** Concentrations of strain JAM1 and strain NL23 in planktonic co-cultures.

| | Concentration<br>gene copies/mL<br>(SD) $\times 10^8$ | Proportion<br>of each<br>strain |
| --- | --- | --- |
| Assays #1 |  |  |
| Strain JAM1 | 6.0 (2.3) | 98.4% |
| Strain NL23* | 0.13 (0.08) | 1.6% |
| Total | 6.1 (2.4) |  |
| Assays #2 |  |  |
| Strain JAM1 | 15.1 (0.9) | 80.3% |
| Strain NL23* | 3.7 (0.3) | 19.7% |
| Total | 18.8 (1.2) |  |

All cultures were performed in the *Methylophaga* 1403 medium with 0.5% NaCl.
\*For the inoculum, strain NL23 was cultured under oxic conditions without $\text{NO}_3^-$ in co-culture assays #1 and under anoxic conditions with 21.4 mM $\text{NO}_3^-$ in co-culture assays #2. qPCR assays were performed with primers targeting *napA* for strain NL23 and *narG1* for strain JAM1. Results are from triplicate cultures. SD: standard deviation.

**Figure 3.**
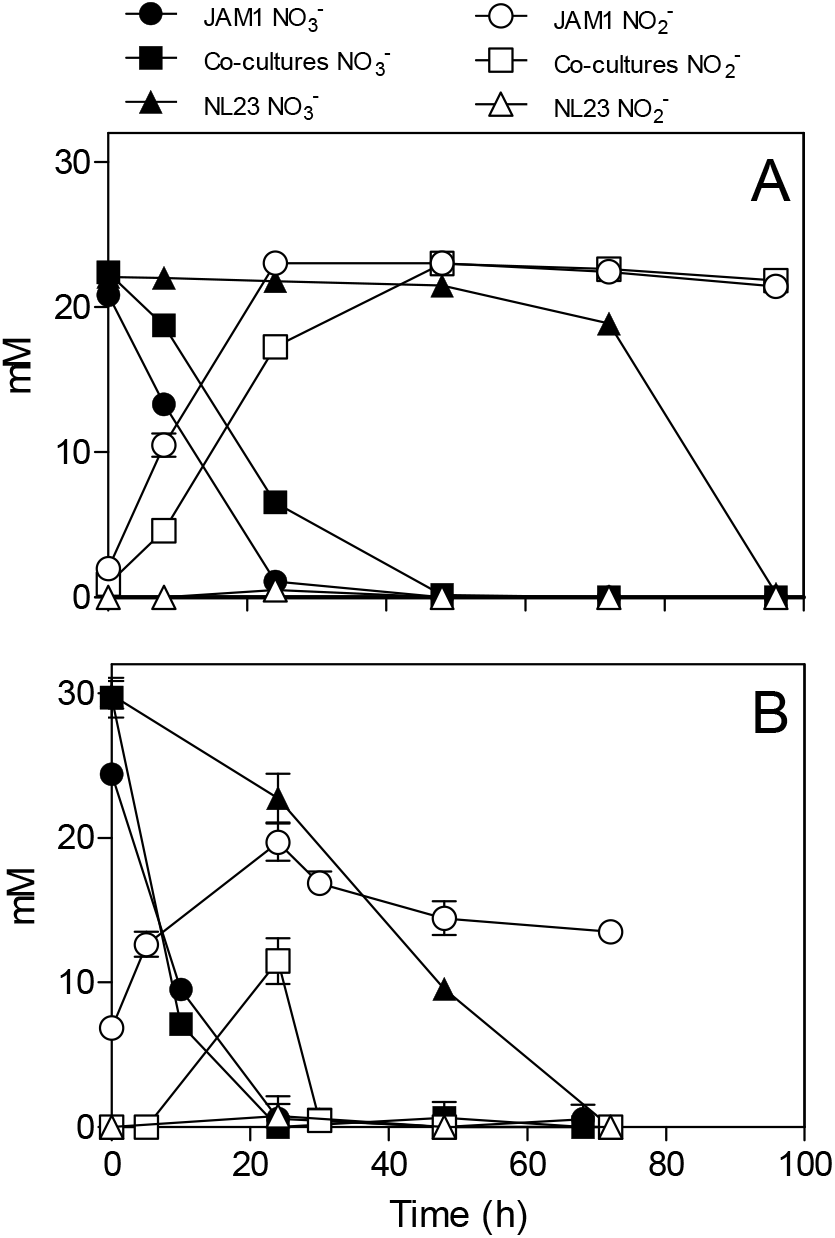
Dynamics of denitrification of the planktonic co-cultures. Strain JAM1 and strain NL23 were co-cultured in the 0.5% NaCl *Methylophaga* 1403 medium under anoxic conditions with 21.4 mM NO_3_− and 0.3% methanol at 30°C. As controls, mono-cultures were carried out with both strains under the same conditions. Concentrations of NO_3_− and NO_2_− were measured. Strain NL23 inoculum was from pre-cultures derived under oxic conditions (no NO_3_−; panel A) or derived under anoxic conditions (with NO_3_−; panel B).

Other planktonic co-cultures were performed by inoculating strain NL23 that was derived from pre-cultures cultured under anoxic conditions to stimulate its denitrification pathway. This time, these co-cultures were capable of full denitrification (Fig. 3B). The NO_3_− reduction rates were similar between the co-cultures (1.165 ± 0.054 mM-NO_3_− h^−1^) and strain JAM1 mono-cultures (0.995 ± 0.032 mM-NO_3_− h^−1^). NO_2_− accumulated transitorily after NO_3_− reduction in the co-cultures and then was completely consumed after 36h (Fig. 3B). The denitrification rates of the co-cultures were twice higher (t-test, p=0.0038) (0.912 ± 0.033 mM-NO_x_ h^−1^) than those in strain NL23 mono-cultures (0.428 ± 0.019 mM-NO_x_ h^−1^). The proportion of strain NL23 was about 12 times higher in these co-cultures than that in the first co-culture assays (19.7% vs 1.6%; Table 4). The total concentrations of both strains in these co-cultures were three times higher than those in the first co-culture assays (Table 4).

### Biofilm co-cultures

As mentioned before, planktonic co-cultures could not be performed under marine conditions without impairing strain NL23 growth. As strain JAM1 and strain NL23 were isolated from a biofilm, developing a biofilm co-culture provided a mean to assess the adaptability of strain NL23 to marine conditions. Two reactors with the Bioflow 9 mm supports and containing the Methylophaga 1403 medium at 0.5% NaCl were inoculated with both strains, and operated under fed-batch mode. The reactors were run for several weeks for the biofilm to build up and the denitrifying activities to stabilize. The biofilm was then acclimated with increasing concentrations of NaCl (Table 1). Finally, the *Methylophaga* 1403 medium was replaced with the Instant Ocean (IO) medium to mimic the original bioprocess. A third reactor was operated under the same conditions, this time with only strain NL23. The three reactors were run sequentially. Results from reactor 1 allowed us to adjust the conditions for reactor 2.

In reactor 1, the NO_3_− reduction rates and the denitrification rates (represented by the NO_x_ reduction rates) ranged both between 1.4 and 5.1 mM h^−1^ (average 2.5 mM h^−1^) when operating with the *Methylophaga* 1403 medium at 0.5%, 1%, 2% and 2.75% NaCl (Table 5). Transitory NO_2_− accumulation was only observed in the reactor operated with 2.75% NaCl, which peaked after 8 h, suggesting a decrease in the denitrification performance. When the reactor was run with the IO medium, an 18-fold decrease occurred in the denitrification rates with accumulation of NO_2_−. The gas production was absent reflecting low bacterial activities. We suspected that the absence of the supplements (Wolf solution, solution T, and B_12_ vitamin) that were present in the *Methylophaga* 1403 medium could have caused this defect. After 12 days following re-addition of these supplements, the activities were partially restored (Table 5).

**Table 5.** Performance of the reactors.

|  | <u>Gas</u><sup>2</sup> | <u>Reduction rates</u><br>mM h<sup>-1</sup> |  | <u>NO<sub>2</sub><sup>-</sup></u><br><u>Peak</u><br>mM (h)<sup>3</sup> | <u>Protein</u><sup>5</sup><br>mg | <u>Specific denitri-</u><br><u>cation rate (NO<sub>x</sub>)</u><br>mM h<sup>-1</sup> mg-prot<sup>-1</sup> | <u>Ectoine</u><br>μg/<br>support |
| --- | --- | --- | --- | --- | --- | --- | --- |
|  | mL | NO<sub>3</sub><sup>-</sup> | NO<sub>x</sub> |  |  |  |  |
| **Reactor 1** Biofilm co-culture |  |  |  |  |  |  |  |
| 0.5%<sup>1</sup> | 85 | 2.01 | 2.01 | none | Nm | Nm | Nm |
| 1.0%<sup>1</sup> | 95 | 1.62 | 1.62 | none | Nm | Nm | Nm |
| 2.0%<sup>1</sup> | 110 | 5.09 | 5.11 | none | Nm | Nm | Nm |
| 2.75%<sup>1</sup> | 100 | 1.40 | 1.44 | 28.4 (8) | Nm | Nm | Nm |
| IO no sup | 0 | 0.25 | 0.08 | 22.4 (96) | Nm | Nm | Nm |
| IO + sup | 45 | 0.86 | 0.10 | 25.1 (24) | 227 | 0.43 | 9.8 |
| **Reactor 2** Biofilm co-culture |  |  |  |  |  |  |  |
| 0.5% | 65 | 2.56 | 0.88 | 6.7 (5) | 217 | 4.05 | 0.6 |
| 1.0% | 110 | 2.73 | 1.30 | 7.1 (4) | 345 | 3.78 | Nm |
| 2.0% | 90 | 5.26 | 1.09 | 13.3 (3) | 183 | 5.94 | 1.4 |
| 2.75% | 110 | 7.14 | 2.19 | 14.1 (2) | 245 | 8.92 | 1.8 |
| IO + sup | 105 | 5.20 | 0.90 | 12.9 (4) | 391 | 1.99 | 17.4 |
| **Reactor 3** Strain NL23 biofilm mono-culture |  |  |  |  |  |  |  |
| 0% | 150 | 5.12 | 5.12 | none | Nm | Nm | Nm |
| 0.5% | 140 | 2.74 | 2.74 | none | 49 | 56.2 | Nm |
| 1.0% | 140 | 2.69 | 2.69 | none | 79 | 34.0 | Nm |
| 2.0% | 90 | 2.70 | 0.53 | 12.9 (8) | 97 | 5.5 | Nm |
| 2.75% | 25 | 3.23 | 0.11 | LC<sup>4</sup> | 34 | 3.3 | Nm |
| IO + sup | 20 | Nm | 0.11 | LC<sup>4</sup> | Nm | Nm | Nm |

<sup>1</sup> Concentration of NaCl in the *Methylophaga* 1403 medium<sup>2</sup> Values are those when the gas production stabilized.<sup>3</sup> Transitory NO<sub>2</sub><sup>-</sup> accumulation: Peak concentration of NO<sub>2</sub><sup>-</sup> (time in hours of the maximum peak measured).<sup>4</sup> NO<sub>2</sub><sup>-</sup> accumulation with low consumption (LC).<sup>5</sup> Estimated total protein in the reactor (suspended biomass and biofilm).**Table 6. Concentrations of the strains JAM1 and NL23 in the reactors determined by qPCR**

In reactor 2, the NO_3_− reduction rates were higher than these measured in reactor 1 when operating with the *Methylophaga* medium at 0.5%, 1%, 2% and 2.75% NaCl (Table 5), and ranged between 2.6 and 7.1 mM h^−1^ (average 4.4 mM h^−1^). These higher NO_3_− reduction rates could have had an impact on the reactor dynamics by generated transitory NO_2_− accumulation that peaked after 2 to 4 hours under these four conditions. Consequently of these transitory accumulations, the denitrification rates in reactor 2 was lower than those in reactor 1 with values ranging between 0.9 and 2.2 mM h^−1^ (average 1.4 mM h^−1^). During these four operating conditions, reactor 2 showed increases from 4.1 to 8.9 mM h^−1^ mg protein^−1^ in the specific denitrification rates. To avoid the detrimental effects observed in reactor 1, reactor 2 was run with the IO medium containing the supplements. The gas production, the NO_3_− reduction and denitrification rates, and the transitory NO_2_− accumulation were similar than those observed in the other operating conditions (Table 5). However, the specific denitrification rates decreased by 4.5 times. Once again, to avoid detrimental effect on the denitrification activities, the supplements were removed one by one (except for trace metal elements) by operating the reactor for a few days between each removal. In all cases, complete denitrification occurred in less than 24 h, and gas production was constant throughout the operating conditions.

In reactor 3 (ran with only strain NL23), the NO_3_− reduction rates ranged between 2.7 and 3.2 mM h^−1^ (average 2.8 mM h^−1^) when operating with the *Methylophaga* 1403 medium at 0.5%, 1%, 2% and 2.75% NaCl (Table 5). However, the denitrification rates and the specific denitrification rates dropped by 5.1 and 6.2 times, respectively, when NaCl concentration reached 2.0%. When operated with 2.75% NaCl and in the IO medium, denitrification rates were further down with NO_2_− accumulation, as well as low gas production.

The proportions of strain NL23 in the suspended biomass of reactor 1 operated with the *Methylophaga* 1403 medium at 0.5%, 1%, 2% and 2.75% NaCl ranged from 2.3% to 21.4% (Table 6). The proportion of strain NL23 was below 1% in reactor 1 operated with the IO medium. The proportions of strain NL23 in the suspended biomass of reactor 2 operated with the *Methylophaga* 1403 medium at 0.5%, 1%, 2% and 2.75% NaCl ranged from 1.3 to 5.9% (Table 6), and were at similar levels in the reactor operated with the IO medium (1.2 and 3.0%). The levels of both strains were also determined in the biofilm of reactor 2. The proportions of strain NL23 were 3-4 times higher in the biofilm (4.3 to 10.9%) than in the suspended biomass, suggesting better environment for strain NL23 to grow. In reactor 3, with the increase of NaCl concentrations in the medium, the concentrations of strain NL23 decreased in both the suspended biomass and the biofilm (Table 6).

**Table 6.** Concentrations of the strains JAM1 and NL23 in the reactors determined by qPCR.

|  | <u>Suspended biomass</u> |  |  | <u>Biofilm</u> |  |  | <u>Total</u><sup>1</sup> | <u>Specific rates</u><sup>2</sup> |
| --- | --- | --- | --- | --- | --- | --- | --- | --- |
|  | <u>cp gene/mL (x10<sup>7</sup>)</u> |  | <u>Proportion</u><br><u>of NL23</u> | <u>cp gene/</u><br><u>support (x10<sup>7</sup>)</u> |  | <u>Proportion</u><br><u>of NL23</u> | <u>cp/</u><br><u>reactor</u> | <u>(μM-NO<sub>x</sub> h<sup>-1</sup>)</u><br><u>per 10<sup>9</sup> gene</u><br><u>copies</u> |
|  | JAM1 | NL23 | % | JAM1 | NL23 | % | (x10<sup>9</sup>) |  |
| **Reactor 1** Biofilm co-culture |  |  |  |  |  |  |  |  |
| 0.5%<sup>1</sup> | 13.2 (28.0) | 0.86 (0.90) | 6.1 | Nm | Nm |  |  |  |
| 1.0% | 1.1 (1.6) | 0.30 (0.11) | 21.4 | Nm | Nm |  |  |  |
| 2.0% | 1.5 (0.5) | 0.21 (0.15) | 12.3 | Nm | Nm |  |  |  |
| 2.75% | 5.0 (1.3) | 0.12 (0.10) | 2.3 | Nm | Nm |  |  |  |
| IO no sup | 42.9 (3.7) | 0.34 (0.2) | 0.78 | Nm | Nm |  |  |  |
| IO + sup | 51.5 (54.6) | 0.32 (0.30) | 0.62 | 56.1 | 8.5 | 1.9 | 453 | 1.9 |
| **Reactor 2** Biofilm co-culture |  |  |  |  |  |  |  |  |
| 0.5% | 60.9 (16.5) | 2.7 (0.96) | 4.2 | 92.8 | 11.4 | 10.9 | 621 | 1.42 |
| 1.0% | 46.2 (12.6) | 2.9 (0.83) | 5.9 | Nm | Nm |  |  |  |
| 2.0% | 23.3 (4.2) | 0.42 (0.15) | 1.8 | Nm | Nm |  |  |  |
| 2.75% | 14.4 (0.9) | 0.19 (0.09) | 1.3 | 262 | 12.8 | 4.7 | 796 | 2.75 |
| IO + sup | 5.2 (4.6) | 0.16 (0.08) | 3.0 | Nm | Nm |  |  |  |
| IO no sup | 10.5 (15.3) | 0.13 (0.17) | 1.2 | 182 | 8.1 | 4.3 | 554 | 1.62 |
| **Reactor 3** Strain NL23 biofilm mono-culture |  |  |  |  |  |  |  |  |
| 0% | NA | 6.7 (3.2) |  | NA | Nm |  |  |  |
| 0.5% | NA | 2.7 (1.4) |  | NA | 23.3 |  | 76 | 36.1 |
| 1.0% | NA | 2.0 (0.86) |  | NA | 15.4 |  | 51 | 52.7 |
| 2.0% | NA | 1.0 (0.59) |  | NA | 8.2 |  | 27 | 19.6 |

An indication of the level of specific denitrifying activities in the reactors was to divide the denitrification rates (mM-NO_x_ h^−1^; in Table 5) by the estimated total number of cells (strain NL23 and strain JAM1) in the reactor (suspended biomass and biofilm) as determined by qPCR (Table 6). These specific denitrification rates, translated per billion of cells (or gene copies), were at similar levels in reactor 2 during the different operating phases (Table 6). In reactor 3, the denitrification rates per billion cells were between 12 to 25 times higher than in reactor 2. It was at its lowest with medium at 2.0% NaCl, suggesting decreases in denitrification efficiency by strain NL23 cells.

### Ectoine in the biofilm co-culture

We measured the level of ectoine in the biofilm of reactor 2 (Table 5). Increasing amount of ectoine was noticed with the increase of NaCl concentration in the reactor.

### Biofilm structure

The colonization of surface by both strains was followed in the reactor with a flow chamber that was connected to the reactor and containing microscope slides (Fig. 1). Figure 4A and B showed uniform dispersed clusters of cells throughout the surface after one to five days of colonization. FISH assays showed that strain NL23 was lower in concentration than strain JAM1 in the biofilm (Fig. 4), which concurs with the qPCR results. No specific pattern of distribution of both strains was noticed in the biofilm.

**Figure 4.**
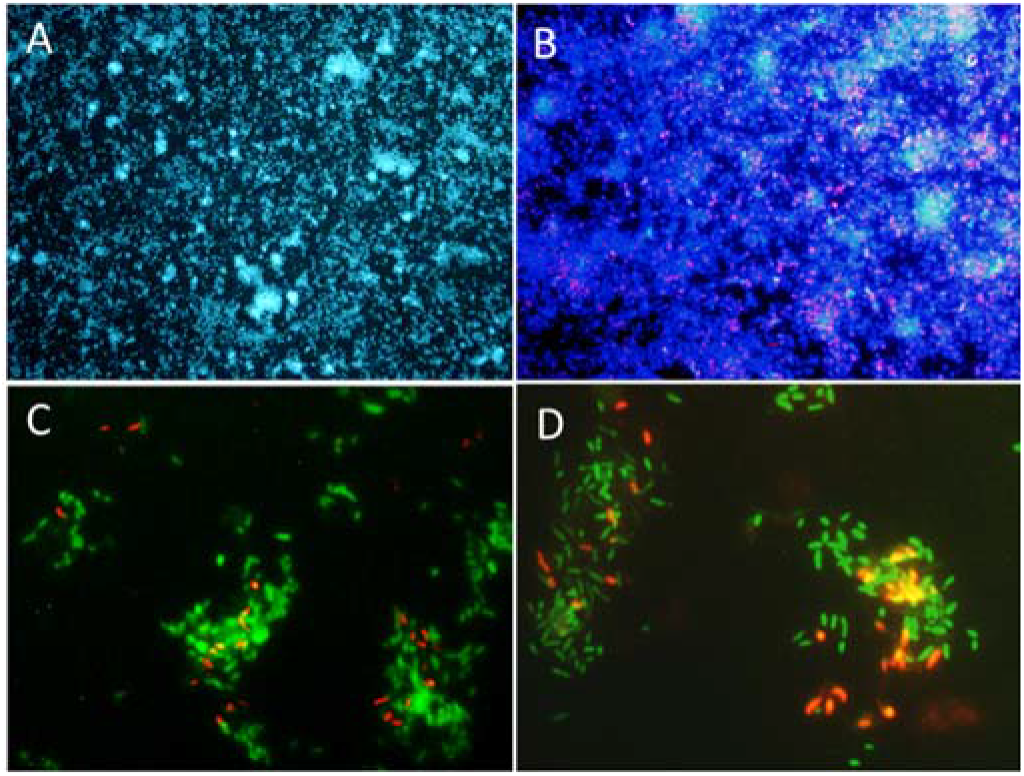
Biofilm examination by fluorescent *in situ* hybridization. Microscope slides were incubated in the flow cell chamber connected to the reactor 2 (biofilm co-culture) for 2-5 days to allow cell colonization. Cells were revealed by DAPI coloration or by FISH with specific probes targeting *Methylophaga* spp. and *Hyphomicrobium* spp., and examined by epifluorescence microscopy. Reactor operated with the *Methylophaga* 1403 medium at: 1% NaCl (panels A, B), at 0.5% NaCl (panel C), and with the IO medium (panel D). A. Total cells revealed by DAPI coloration (200X). B. *H. nitrativorans* NL23 (magenta) overlay with DAPI coloration (200X). C-D. *H. nitrativorans* NL23 (orange) and *M. nitratireducenticrescens* JAM1 (green) (1000X).

### Gene expression of denitrification genes in biofilm co-culture and mono-culture

The impact of the operating conditions in the reactors on the expression of denitrification genes of key reductases were assessed by measuring in the biofilm the levels of transcripts of both *nar* genes (*narG1* and *narG2*) for strain JAM1, and *napA* and *nirK* for strain NL23. The transcript levels of *narG1* did not change in the biofilm co-culture of reactor 2 operated at 0.5% and 2.75% NaCl, and with the IO medium (Table 7). For *narG2*, the transcript levels in the biofilm co-culture raised by about 2-3 times when the operating conditions in reactor 2 increased in NaCl concentration from 0.5% to 2.75% or when the medium was changed for the IO medium (Table 7).

**Table 7.** Changes in the transcript levels of selected denitrification genes.

|  | Strain NL23 |  | Strain JAM1 |  |
| --- | --- | --- | --- | --- |
|  | *napA* | *nirK* | *narG1* | *narG2* |
| Reactor 2 |  |  |  |  |
| 0.5% | 630 (347)<sup>a</sup> | 478 (219)<sup>a</sup> | 151 (18)<sup>a</sup> | 54 (5)<sup>a</sup> |
| 2.75% | 56 (18)<sup>b</sup> | 224 (154)<sup>ab</sup> | 185 (33)<sup>a</sup> | 88 (12)<sup>b</sup> |
| IO | 37 (6)<sup>b</sup> | 77 (31)<sup>b</sup> | 182 (100)<sup>a</sup> | 140 (42)<sup>c</sup> |
| Reactor 3 |  |  |  |  |
| 0.5% | 40 (19)<sup>b</sup> | 180 (143)<sup>ab</sup> | NA | NA |
| 2.0% | 15 (7)<sup>b</sup> | 286 (155)<sup>ab</sup> | NA | NA |

Values are expressed as gene copies per 100 copies of *rpoB*. Values under parentheses are standard error. Superscript letter: For a specific gene, values with the same superscript letter between the biofilm samples are not significantly different (One-way analysis of variance, $P < 0.05$ , Tukey's multiple comparison test). NA: not applicable.

The *napA* transcript levels in the biofilm co-culture of reactor 2 had an 11-fold decrease when the operating conditions changed from 0.5% NaCl to 2.75% NaCl, or a 17-fold decrease when the medium was changed for the IO medium (Table 7). The *napA* transcript levels were similar in the biofilm mono-culture of reactor 3 operated with 0.5% and 2.0% NaCl. Finally, the *nirK* transcript levels showed no significant difference between the operating conditions in reactors 2 and 3, except in the biofilm co-culture of reactor 2 operated with the IO medium with a 6-fold decrease compared to the biofilm co-culture of reactor 2 operated with 0.5% NaCl.

### The transcriptomic analysis of selected biofilm

The impact of the operating conditions in gene expression of the pathways of denitrification and nitrogen assimilation were assessed by deriving the transcriptome of biofilm samples taken from reactors B2 and B3 operated in different conditions (Fig. 5). No substantial changes were found in the relative transcript levels of the denitrification genes between the three operating conditions in reactor B2 (0.5% and 2.75% NaCl and IO) for both strains, and in the two operating conditions in reactor B3 (0.5% and 2.0% NaCl) for strain NL23.

**Figure 5.**
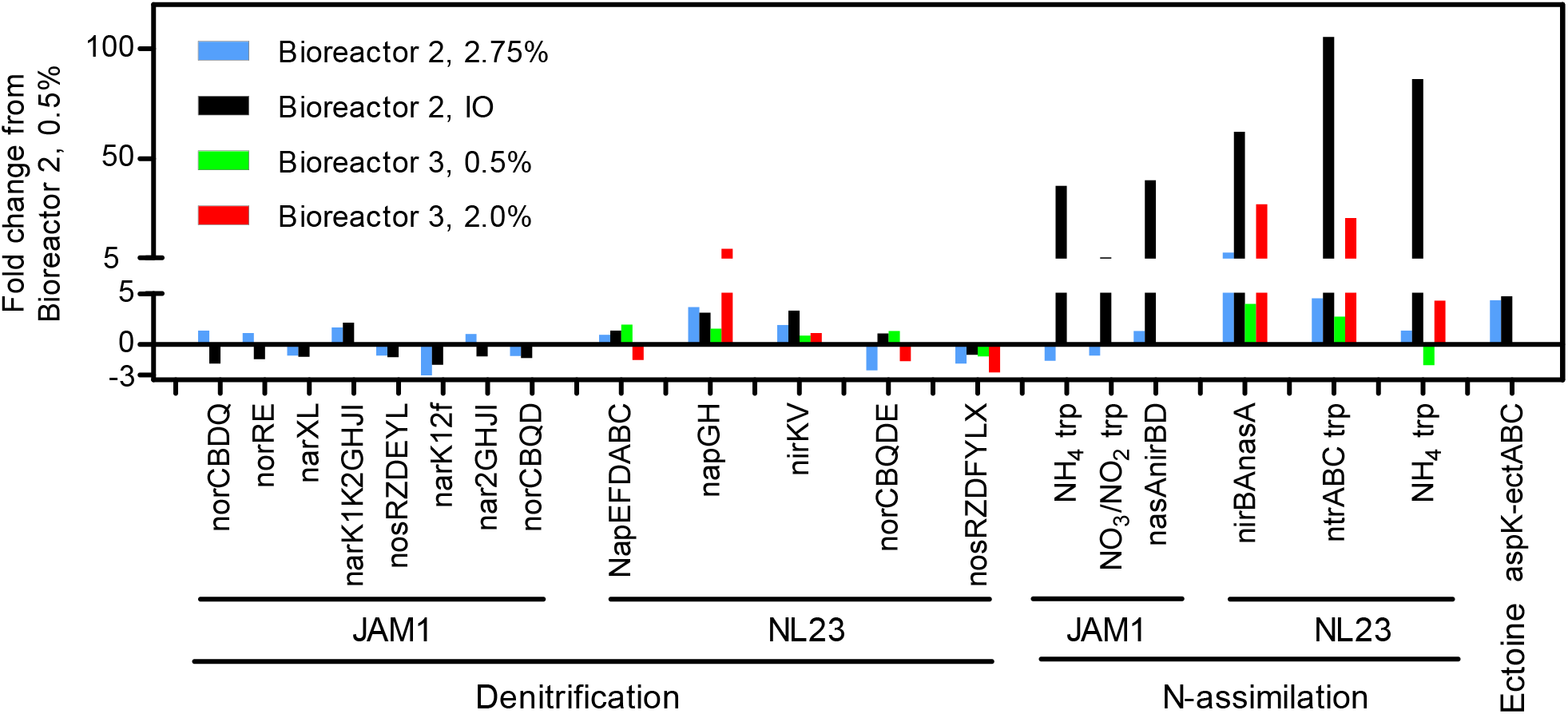
Changes in the relative transcript levels of genes involved in the denitrification and N-assimilation pathways, and in ectoine synthesis. Total RNA were extracted from the biofilm samples taken from the bioreactors 2 and 3 operated at the different conditions. RNA samples were sequenced, and the reads associated to the denitrification genes, the N-assimilation genes and ecotine genes were transformed in transcripts per million (TPM). The TPM values of each gene operon were compared to those of the bioreactor 2, 0.5% NaCl, which was set to 1.0.

The type of medium (*Methylophaga* 1403 medium and IO medium) had a great impact on the N-assimilation pathways in both strains. Higher relative transcript levels were observed in both strains in the reactor B2 and B3 that were running with the IO medium for genes encoding ammonium transporters, and genes involved in the NO_3_− transformation in ammonium. The *Methylophaga* 1403 medium contains ammonium, which is absent in the IO medium. Therefore, the biofilm cultured with the IO medium relied on N-assimilation pathways as nitrogen source.

Finally, the relative transcript levels of the four genes involved in the ectoine synthesis in strain JAM1 were higher in reactor 2 operated under marine conditions (2.75% NaCl and IO medium) compared to the reactor operated in low salt concentration (0.5% NaCl).

## DISCUSSION

The marine and methylotrophic environment that prevailed in the original denitrification system favored the establishment in the biofilm of two methylotrophic species, *H. nitrativorans* and *M. nitratireducenticrescens* directly involved in the denitrifying activities. An equilibrium between these two species had to be reached in this biofilm where specific physiological characters of both species may have certainly played important roles in this equilibrium. One of these characters is the dynamics of NO_3_− uptake. In pure cultures, *M. nitratireducenticrescens* strain JAM1 has shown to have a readiness to reduce NO_3_− with no lag phase for growth contrary to *H. nitrativorans* strain NL23, which has a 2-3 days lag phase before NO_3_− starts to be consumed and growth to occur. These results suggest that the denitrification pathway is tightly regulated in strain NL23 under anoxic conditions, which is not in strain JAM1. Compared to strain NL23, strain JAM1 has a higher μmax for growth and higher specific NO_3_− reduction rates, suggesting that strain JAM1 cells have a higher dynamism of NO_3_− processing (*e.g.* NO_3_− intake and reduction) than strain NL23 cells. These dynamisms might be attributed to the different types of NO_3_− reductases used by these strains. Strain JAM1 possesses two Nar systems, which contribute both to the growth of strain JAM1 under anoxic conditions [35], whereas strain NL23 has one periplasmic Nap-type NO_3_− reductases [12]. The Nar system is found associated with the membrane facing the cytoplasm. NO_3_− reduction by this system can generate proton motive force (PMF) and thus energy for the cells.

Because of its location, the Nap system does not generate PMF upon NO_3_− reduction [3]. Furthermore, strain JAM1 possesses three NO_3_− transporters associated with the denitrification pathway. As the antagonist assays did not show exclusion from each others in the tested conditions, differences in the regulation and the kinetics of the denitrification pathway could therefore confer an advantage of strain JAM1 for growth over strain NL23. These features are reflected in the higher abundance of strain JAM1 than strain NL23 that we found in the co-cultures as revealed by qPCR and FISH assays.

The planktonic co-cultures assays revealed the importance of strain NL23 to have its denitrification pathway stimulated to compete with strain JAM1 for growth. This stimulation resulted in better denitrification rates in planktonic co-cultures compared to those measured in strain NL23 pure cultures. As the original denitrification system operated under continuous mode, with constant concentration of NO_3_− in the affluent, strain NL23 would have had its denitrification pathway stimulated, thus capable to sustain growth.

The planktonic co-cultures did not reflect the true co-habitation of both strains under marine conditions that prevailed in the original denitrification system, since the planktonic co-cultures were carried out at low NaCl concentration to allow strain NL23 to grow. Our hypothesis that strain JAM1 has to be present for strain NL23 to survive under marine conditions revealed to be true with the biofilm co-culture assays in the reactors. Acclimation of strain NL23 with increasing concentration of NaCl and then to the marine IO medium (medium used in the aquarium tanks that were treated by the original system) showed (i) sustained denitrifying activities, (ii) support colonization of strain by NL23 (FISH assays) and (iii) higher proportions of strain NL23 in the biofilm co-culture compared to the suspended biomass.

With the increase of NaCl concentration in reactor 3 (ran only with strain NL23), important decreases in denitrifying activities was noticed, specially when the NaCl concentration reached 2.0%. Martineau *et al.* [12] showed decreases in growth and denitrifying activities in pure cultures of strain NL23 cultured at 2% NaCl. However, compared to reactor 2 (co-cultures), reactor 3 had higher specific denitrification rates (at least in low NaCl concentrations) and higher denitrification rates per billions of cells. These differences in favor of strain NL23 can be explained by the lower level of biomass generated in this reactor (about 4 times less in protein; Table 5). Therefore, the presence of strain JAM1 did not contribute in better specific denitrifying activities in the reactor. However, its presence was essential for strain NL23 to operate at NaCl concentrations higher than 1.0%.

No substantial changes in the transcript levels of the measured denitrification genes in the biofilm co-culture of reactor 2 were noticed except for *napA* when the reactor was operated at 0.5% NaCl. In these conditions, the *napA* transcript levels were >10 times higher than those measured in the other operating conditions (2.75% NaCl and IO medium). However, such levels of *napA* transcripts were not observed in the biofilm mono-culture of reactor 3 operated with 0.5% NaCl, suggesting that strain JAM1 could have had an impact on the *napA* expression in the biofilm co-culture at this NaCl concentration. Wang *et al.* [38] showed decrease of about 10 times in denitrification activities of a denitrifying granular sludge when the NaCl concentration increased gradually from 0% to 10% in synthetic wastewater. The same study observed increases in lethality and DNA leakage in these cultures when the NaCl concentrations were above 2%. Not many studies were reported on the effect of salinity on the expression of denitrification genes. Gui *et al.* [39] observed decreases in the expression of the 4 denitrification genes (including *napA*) with the increase of NaCl concentration in the medium of *Achromobacter* sp. GAD-3 pure cultures.

We observed a drop of the denitrifying activities with the passage from the *Methylophaga* 1403 medium to the IO medium. Besides IO to be commercial medium which could have unknown additives, differences between the two media is the presence of phosphate, ammonium and BisTris (buffer, and can bind some ions) in the *Methylophaga* 1403 medium that are absent in the IO medium. Ammonium can be a source of N that is readily assimilated in the glutamine pathway for biomolecule synthesis. In IO medium, nitrate must be transformed in ammonium before this assimilation, which require energy from the cells. Indeed, we found higher levels of relative transcripts of genes associated with the N-assimilation pathways of both strains in the biofilms that were derived from reactors operated with IO.

As expected, increase in ectoine production in the biofilm co-culture correlated with the increase in NaCl concentration in reactor 2. This higher level concurs with the higher relative transcript levels of the genes involved in ectoine production in the biofilm derived from the reactor 2 operated with 2.75% NaCl and IO compared to this reactor operated with 0.5% NaCl. Acclimation by strain NL23 of increases of NaCl concentration in the biofilm co-cultures could be related with an increase in ectoine production by strain JAM1. It is possible that the level of strain NL23 that can be reached in the biofilm is restricted by the level of ectoine generated by strain JAM1. Many bacteria have developed uptake transporter systems to accumulate osmoprotectant such as ectoine in their cells. For instance, *Corynebacterium glutamicum* possesses four secondary carriers for compatible solutes among which two, ProP and EctP, can uptake ectoine. These transporters belong respectively to the major facilitator superfamily and to the sodium/solute symporter superfamily [40]. Genes associated with these types of transporters were found in the genome of strain NL23 [13]. Huang ZZ *et al.* [41] found high levels of intracellular ectoine and hydroxyectoine in a phenol-degrading microbial enrichment acclimated up to 3 M NaCl (17.5%). Demonstration that osmoproctectant uptake by bacteria in biofilm occurs was shown by Kapfhammer *et al.* [42] where a *Vibrio cholera* strain impaired in the production of ectoine can increase cell attachment and biofilm growth in presence of the osmoprotectant glycine betaine in high-osmolarity medium, but not when the betaine transporter was impaired too.

## CONCLUSIONS

*M. nitratireducenticrescens* strain JAM1 have a higher dynamism of NO - processing (*e.g.* NO_3_− intake and reduction) than *H. nitrativorans* strain NL23. Differences in the regulation and the kinetics of the denitrification pathway confer an advantage to strain JAM1 for growth over strain NL23. Better denitrifications rates in planktonic co-cultures were obtained in low salt medium. In reactors carrying the biofilm co-culture, acclimation of strain NL23 with increasing concentration of NaCl and then in the marine IO medium showed (i) sustained denitrifying activities, (ii) support colonization by strain NL23 and (iii) higher proportions of strain NL23 in the biofilm compared to the suspended biomass. Reactor ran with only strain NL23 failed to achieve such activities under marine conditions. Higher concentrations of ectoine were measured in the biofilm co-culture with the reactor operated under marine conditions. Although the presence of strain JAM1 did not contribute in better specific denitrifying activities in the reactor, its presence was essential for strain NL23 to operate at NaCl concentrations > 1.0%. Our results highlighted some mechanisms of co-habitation from these two strains in achieving denitrifying activities, and can contribution in optimizing of denitrification systems treating saline/brackish waters.

## ACKNOWLEDGEMENTS

We thank Karla Vasquez and Sylvain Milot for her technical assistance.

